## Supplemental Information for "On the effect of phylogenetic correlations in coevolution-based contact prediction in proteins"

The supporting information contains the supplementary tables and figures.

| Pfam ID | L | M | PDB ID |
|---|---|---|---|
| PF02906 | 243 | 4549 | 1feh |
| PF11976 | 72 | 2467 | 5gjl |
| PF00786 | 59 | 2747 | 1f3m |
| PF00988 | 128 | 8611 | 5dot |
| PF00338 | 98 | 6983 | 2mew |
| PF00617 | 177 | 7173 | 3t6a |
| PF02787 | 122 | 9080 | 1a9x |
| PF02609 | 52 | 6068 | 1vp7 |
| PF10369 | 74 | 6405 | 2f1f |

**Table 1.** Protein families and PDB structures used in dataset DS1. The alignment depth $M$ is counted after removal of duplicated sequences.

| Pfam ID | L | M | PDB ID | Pfam ID | L | M | PDB ID |
|---|---|---|---|---|---|---|---|
| PF00301 | 47 | 4051 | 1brf | PF00014 | 53 | 10628 | 1aap |
| PF00048 | 60 | 4148 | 1m8a | PF01363 | 69 | 11022 | 1vfy |
| PF00542 | 67 | 7670 | 1ctf | PF07647 | 66 | 7062 | 1kw4 |
| PF05198 | 70 | 6442 | 1tif | PF00036 | 29 | 5487 | 1avs |
| PF00051 | 79 | 4526 | 1i71 | PF02214 | 94 | 7731 | 1dsx |
| PF10150 | 270 | 7777 | 5f6c | PF00030 | 82 | 8121 | 1nps |
| PF00381 | 82 | 9321 | 1pch | PF00234 | 87 | 1871 | 1fk5 |
| PF04296 | 75 | 3552 | 1g2r | PF01985 | 84 | 5759 | 1jo0 |
| PF00127 | 99 | 3946 | 1ag6 | PF00077 | 101 | 1313 | 2hs1 |
| PF02033 | 105 | 6787 | 1jos | PF01807 | 98 | 8001 | 1doq |
| PF02302 | 90 | 9193 | 1iib | PF07653 | 57 | 6654 | 1i1j |
| PF01250 | 90 | 7316 | 1vmb | PF00237 | 103 | 8664 | 1i4j |
| PF00031 | 92 | 3341 | 1roa | PF01668 | 143 | 6633 | 1wjx |
| PF03960 | 64 | 8699 | 1rw1 | PF01152 | 121 | 5111 | 1dlw |
| PF00238 | 122 | 7535 | 1whi | PF00116 | 120 | 7691 | 2cua |
| PF02036 | 101 | 6687 | 1c44 | PF02579 | 94 | 3530 | 1p9o |
| PF00572 | 119 | 8938 | 1j3a | PF00241 | 125 | 7343 | 1m4j |
| PF00565 | 108 | 9233 | 1ihz | PF01389 | 183 | 594 | 1qjp |
| PF00080 | 141 | 6259 | 1ej8 | PF01687 | 121 | 7932 | 1nb9 |
| PF00042 | 110 | 7336 | 1a6m | PF06445 | 155 | 6600 | 1jyh |
| PF00061 | 144 | 4886 | 1beb | PF00413 | 158 | 6350 | 1hfc |
| PF02367 | 128 | 6576 | 1htw | PF00959 | 109 | 2509 | 1lpy |
| PF01812 | 187 | 8531 | 1wkc | PF02233 | 460 | 4345 | 1d4o |
| PF01195 | 183 | 8170 | 1ryb | PF00186 | 160 | 7083 | 1aoe |
| PF01661 | 118 | 8797 | 1vhu | PF01339 | 170 | 7750 | 1chd |
| PF01421 | 200 | 6099 | 1atl | PF04509 | 38 | 3446 | 1xkra |
| PF00445 | 185 | 4081 | 1dix | PF00045 | 45 | 8482 | 1hxn |
| PF02224 | 211 | 6974 | 1cke | PF01183 | 178 | 5228 | 1jfx |
| PF02527 | 184 | 6748 | 1xdz | PF00310 | 420 | 7072 | 1xff |
| PF00139 | 249 | 5676 | 1gzc | PF01223 | 226 | 5147 | 1ql0 |

**Table 2.** Dataset DS2: Protein families and PDB structures extracted from the PSICOV benchmark, with MSA sizes obtained from current databases. The alignment depth $M$ is counted after removal of duplicated sequences. The selection criterion was to choose those proteins whose MSAs after removing the duplicate sequences did not exceed 12000 sequences, this led to the 60 proteins presented.

| Pfam ID | L | M | PDB ID | Pfam ID | L | M | PDB ID |
|---|---|---|---|---|---|---|---|
| PF02829 | 97 | 967 | 1j5y | PF01896 | 182 | 1770 | 4bpu |
| PF12397 | 116 | 950 | 5wy4 | PF14720 | 80 | 1593 | 3myr |
| PF06003 | 264 | 1394 | 4gli | PF03592 | 141 | 1072 | 3zqn |
| PF04006 | 617 | 1570 | 6nd4 | PF02888 | 75 | 489 | 6cno |
| PF16677 | 120 | 185 | 3p9a | PF17842 | 147 | 142 | 3htx |
| PF18441 | 136 | 131 | 3htx | PF11053 | 157 | 112 | 3txs |
| PF08468 | 157 | 611 | 2pjd | PF02723 | 75 | 54 | 5x29 |
| PF02380 | 93 | 71 | 2pf4 | PF02315 | 88 | 81 | 5xm3 |
| PF15127 | 92 | 109 | hvz | PF02287 | 133 | 241 | 1dio |
| PF16785 | 111 | 145 | 3u8v | PF08640 | 83 | 985 | 5ic8 |

**Table 3.** Dataset DS3: Protein families and PDB structures with small MSA from current databases. The alignment depth $M$ is counted after removal of duplicated sequences. The selection criterion was to choose Pfam families whose alignments depth varies between 50-1800 sequences and whose structures are well known, this led to the 20 families presented.

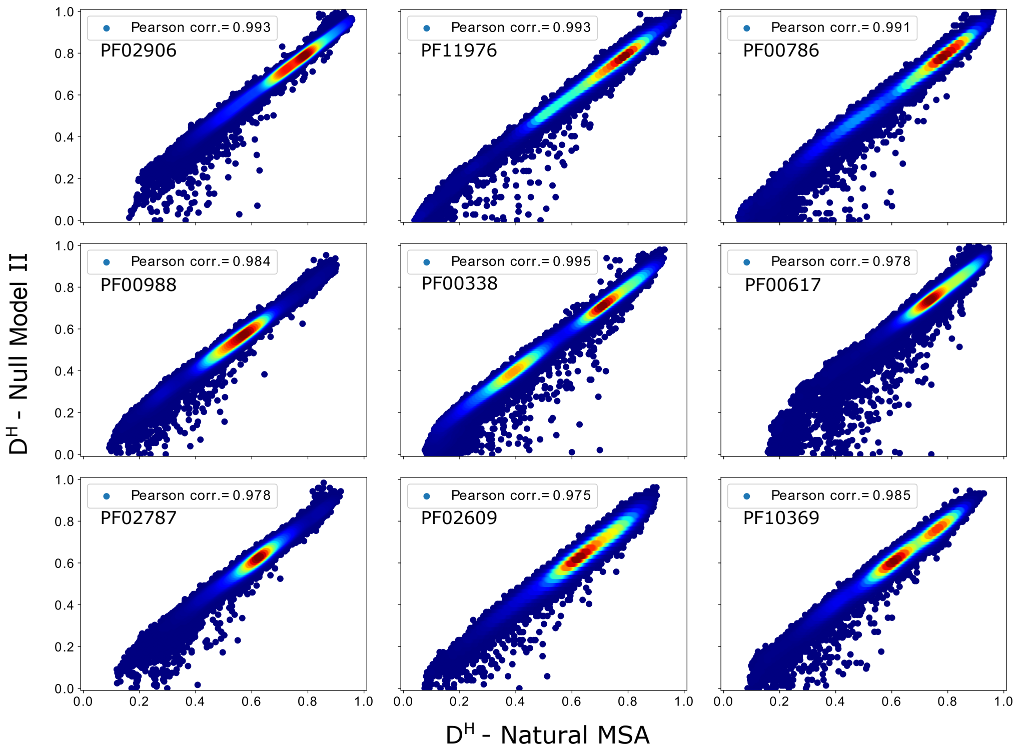


**Fig 1. Scatter plots of pairwise distances between sequences** in the natural MSA vs. an MSA generated by Null model II, for all families in dataset DS1. The inserts show the Pearson correlation between the two distance matrices.

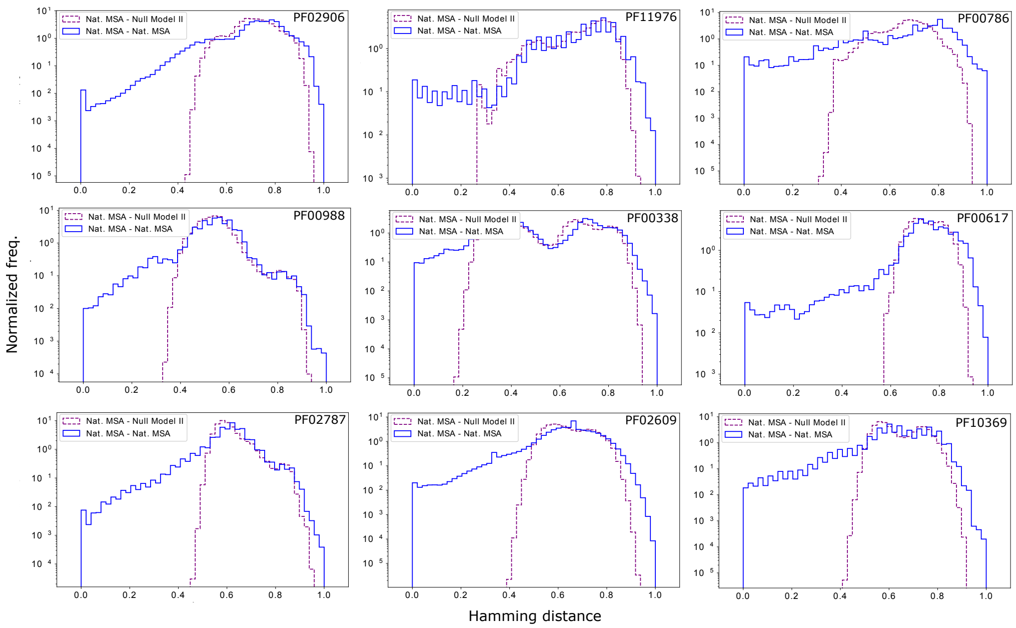


**Fig 2. Histograms of pairwise distances between sequences** in the natural MSA vs. an MSA generated by Null models I and II, for all families in dataset DS1.

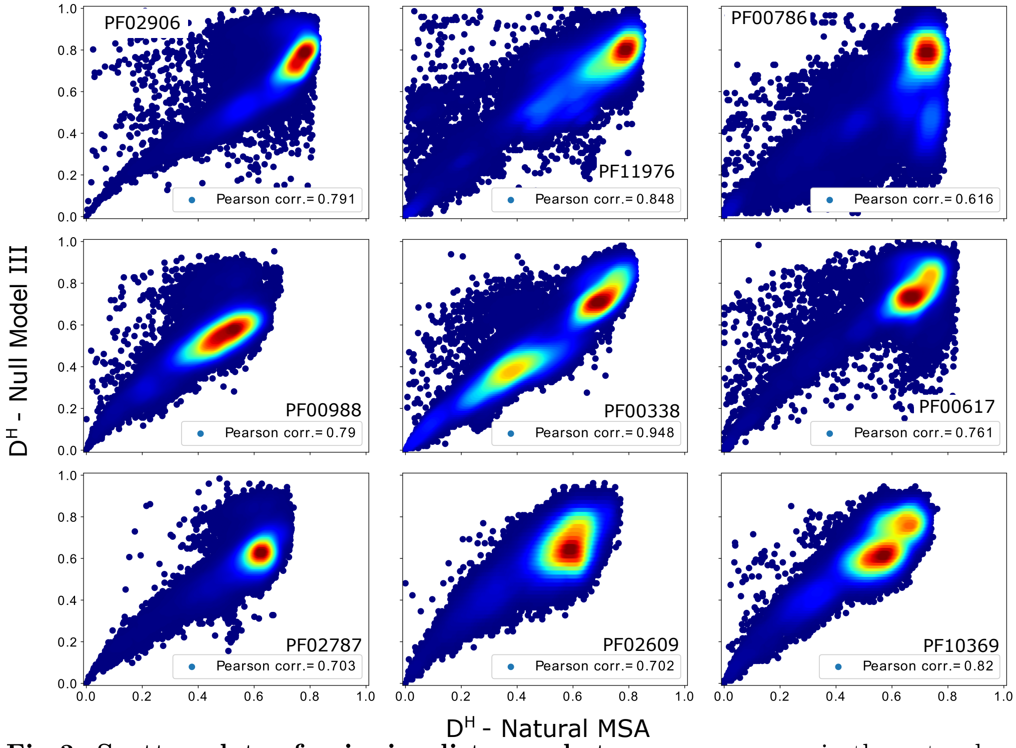


**Fig 3. Scatter plots of pairwise distances between sequences** in the natural MSA vs. an MSA generated by Null model III, for all families in dataset DS1. The inserts show the Pearson correlation between the two distance matrices.

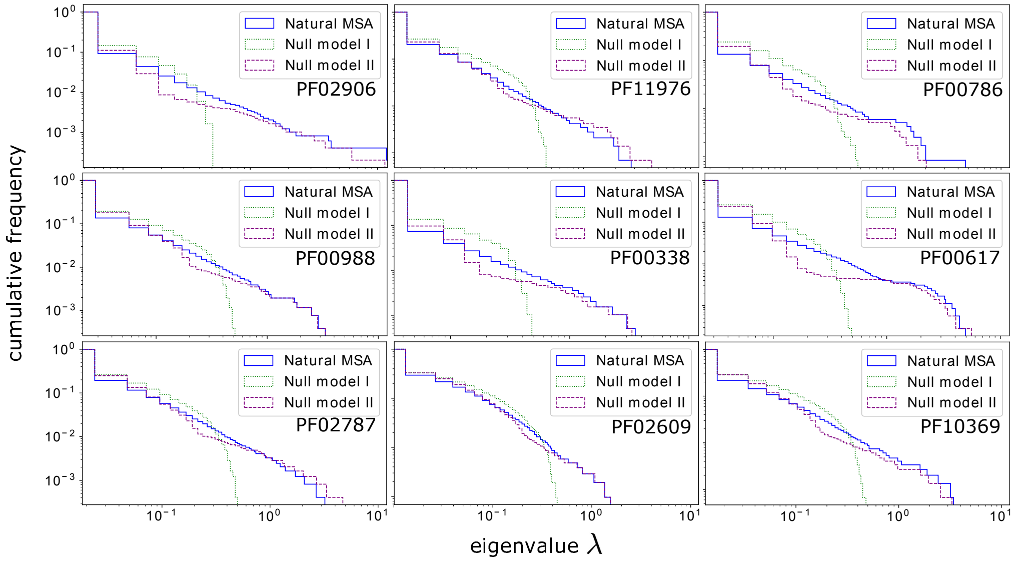


**Fig 4. Eigenvalue spectra of the covariance matrix of the natural MSA and for Null models I and II:** We show cumulative distributions of the eigenvalue spectra for the nine protein families in DS1, i.e. the fraction of eigenvalues larger than $\lambda$ is shown as a function of $\lambda$. We observe that the phylogeny-aware Null model II shows the same fat tail for large eigenvalues, which is also present in the natural data, while the non-phylogenetic Null model I has a more compact support. Data for the Null models are averaged over 50 independent realizations each.

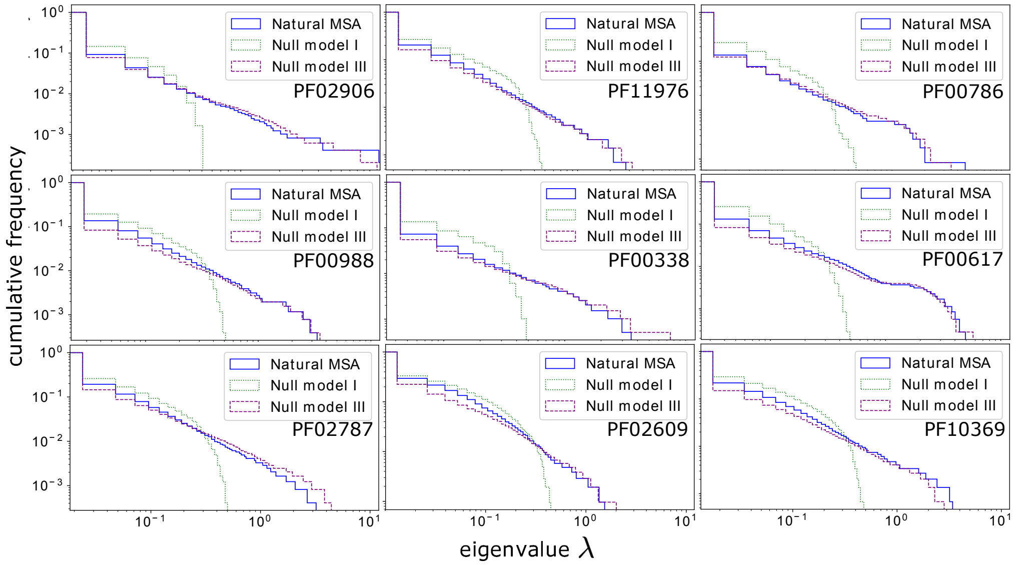


**Fig 5. Eigenvalue spectra of the covariance matrix of the natural MSA and for Null models I and III:** We show cumulative distributions of the eigenvalue spectra for the 9 protein families in DS1, i.e. the fraction of eigenvalues larger than $\lambda$ is shown as a function of $\lambda$. We observe that the phylogeny-aware Null model III shows the same (or an even larger) fat tail for large eigenvalues, which is also present in the natural data, while the non-phylogenetic Null model I has a more compact support. Data for the Null models are averaged over 50 independent realizations each.

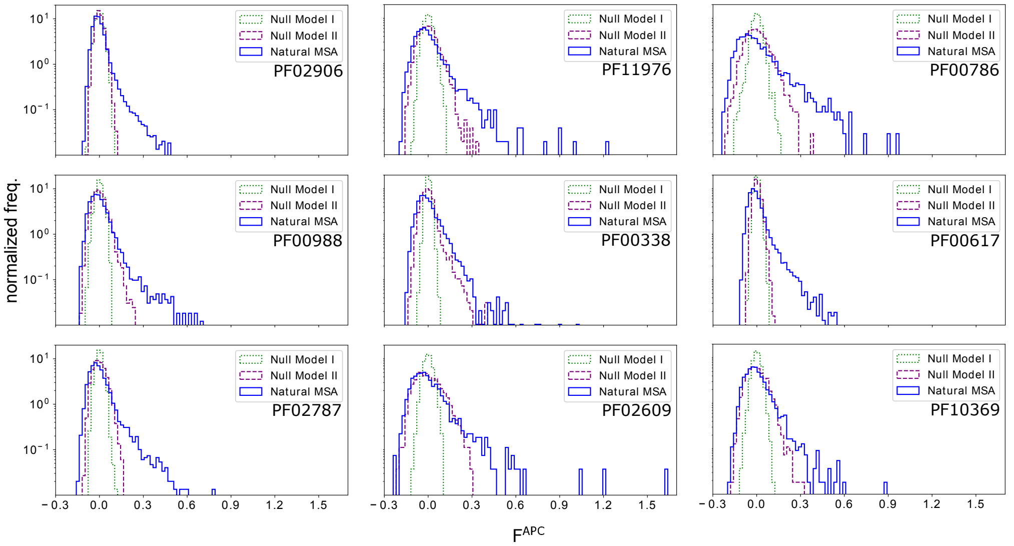


**Fig 6. Histogram of DCA scores derived from natural sequence data and from MSA generated by Null models I and II :** For the 9 protein families in DS1, we show the histograms of DCA coupling scores (APC corrected Frobenius norm of couplings, the standard output of plmDCA), for the natural MSA and samples of Null models I and III. It becomes evident that phylogenetic effects create, to a degree varying from family to family, larger couplings than to be expected from finite sample size alone. However, couplings derived from the natural MSA have substantially larger values.

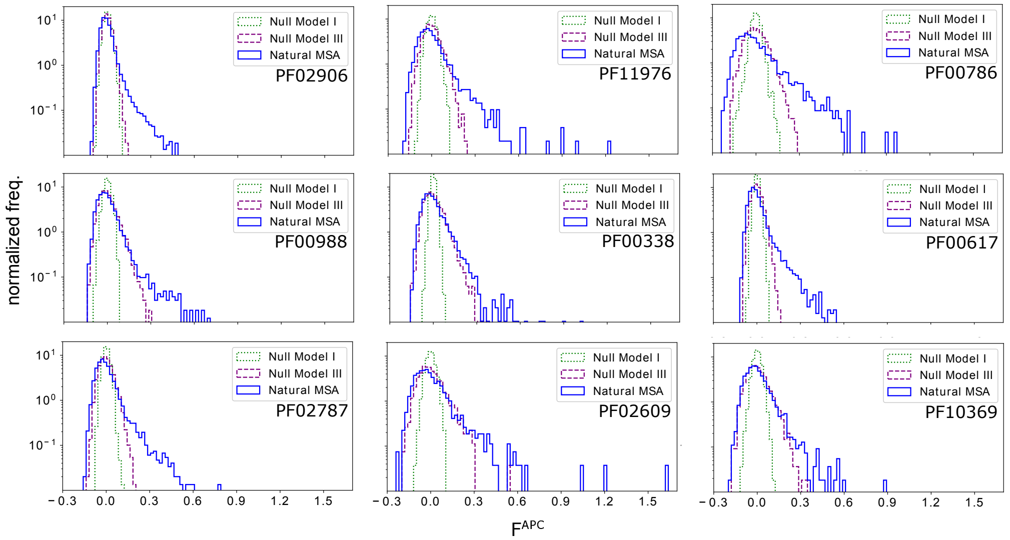


**Fig 7. Histogram of DCA scores derived from natural sequence data and from MSA generated by Null models I and III :** For the protein families in dataset DS1, we show the histograms of DCA coupling scores (APC corrected Frobenius norm of couplings, the standard output of plmDCA), for the natural MSA and samples of Null models I and III. It becomes evident that phylogenetic effects create, to a degree varying from family to family, larger couplings than to be expected from finite sample size alone. However, couplings derived from the natural MSA have substantially larger values.

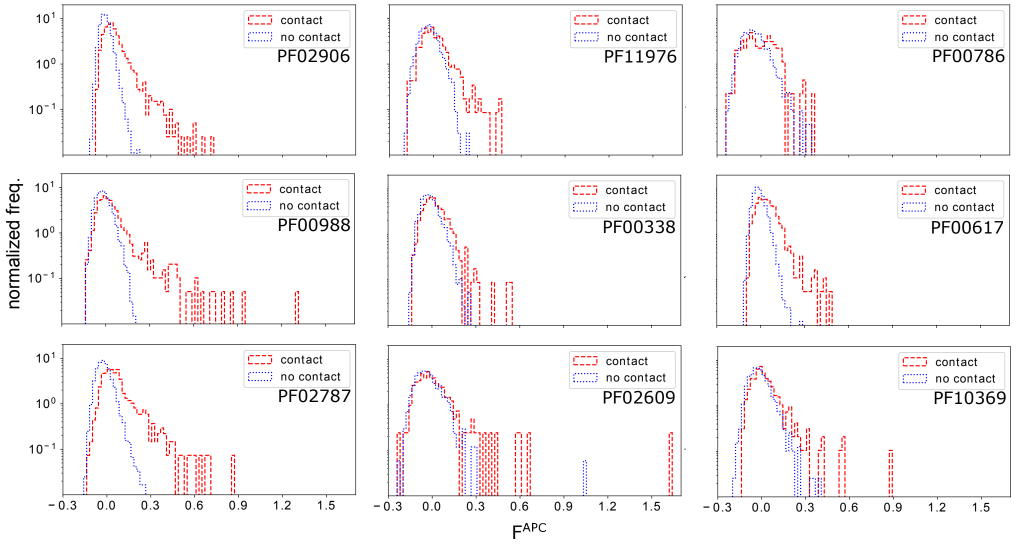


**Fig 8. Histogram of DCA scores derived from natural sequence data for residue-residue contacts and non-contacts:** For the protein families in dataset DS1, we show the histograms of DCA coupling scores (APC corrected Frobenius norm of couplings, the standard output of plmDCA), separated for contacts and non-contacts. Only pairs with linear separation $|i - j| > 4$ along the chain are taken into account. It is clear that for scores above about 0.2-0.3, most predictions are true contacts (false positives may actually be oligomeric contacts), and the quality of contact prediction mostly depends on how many predictions above this threshold are found in a protein family.

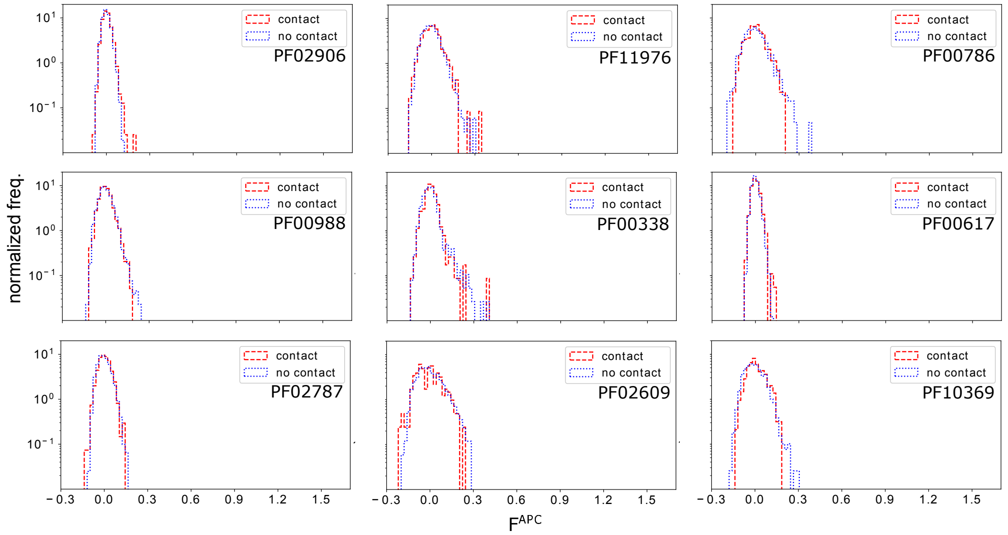


**Fig 9. Histogram of DCA scores derived from Null model II for residue-residue contacts and non-contacts:** For the protein families in dataset DS1, we show the histograms of DCA coupling scores (APC corrected Frobenius norm of couplings, the standard output of plmDCA), separated for contacts and non-contacts. Only pairs with linear separation $|i - j| > 4$ along the chain are taken into account. It becomes evident that any signal related to contacts is totally destroyed by the randomization procedure in Null model II. Interestingly, the null model generates almost no couplings with scores above 0.2-0.3, which was seen in Fig. 8 as a cutoff for high-accuracy contact prediction.

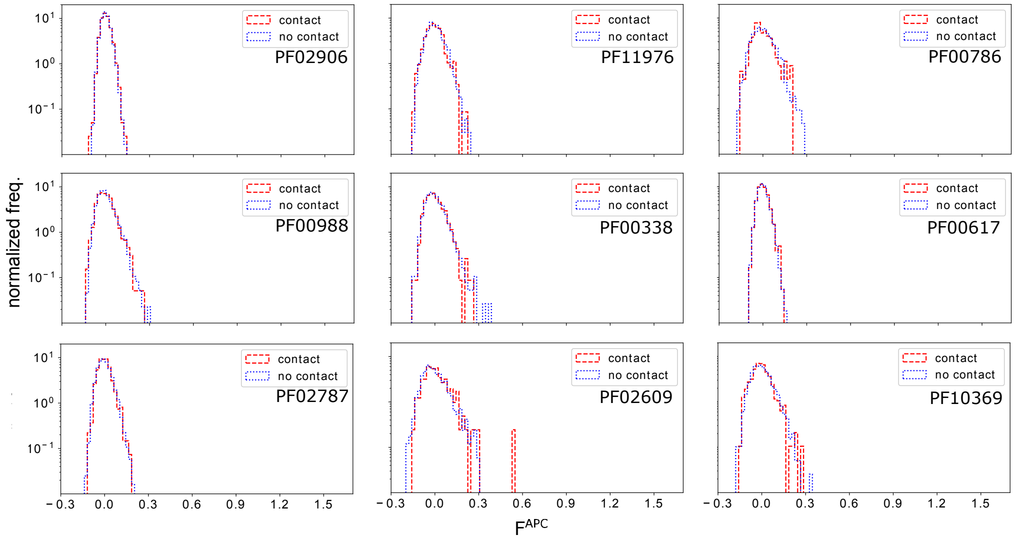


**Fig 10. Histogram of DCA scores derived from Null model III for residue-residue contacts and non-contacts:** For the protein families in dataset DS1, we show the histograms of DCA coupling scores (APC corrected Frobenius norm of couplings, the standard output of plmDCA), separated for contacts and non-contacts. Only pairs with linear separation $|i - j| > 4$ along the chain are taken into account. It becomes evident that any signal related to contacts is totally destroyed by the randomization procedure in Null model III. Interestingly, the null model generates almost no couplings with scores above 0.2-0.3, which was seen in Fig. 8 as a cutoff for high-accuracy contact prediction.

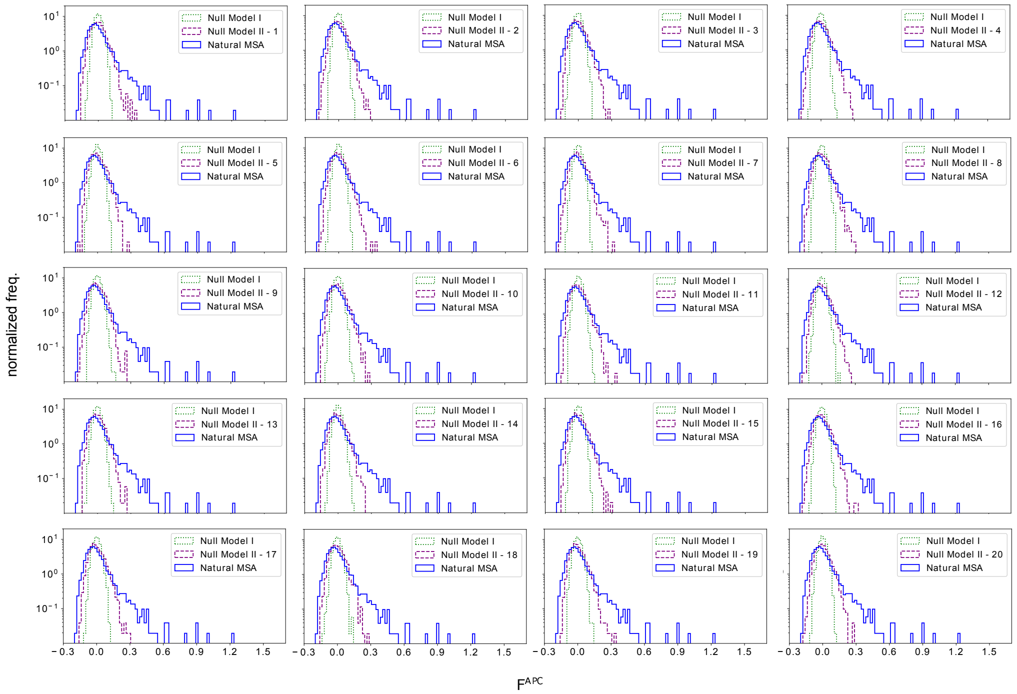


**Fig 11. Histograms of DCA scores derived from 20 independent samples of Null model II for protein family PF02906:** The histograms of couplings are robust with respect to sample-to-sample fluctuations of Null model II.

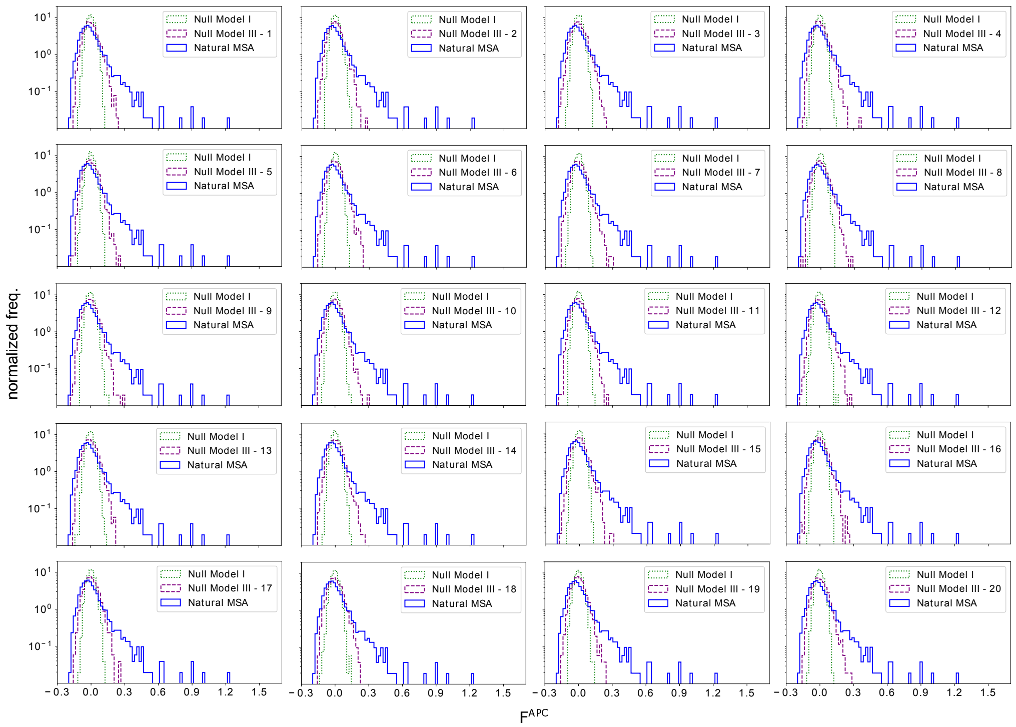

**Fig 12. Histograms of DCA scores derived from 20 independent samples of Null model III for protein family PF02906:** The histograms of couplings are robust with respect to sample-to-sample fluctuations of Null model III.

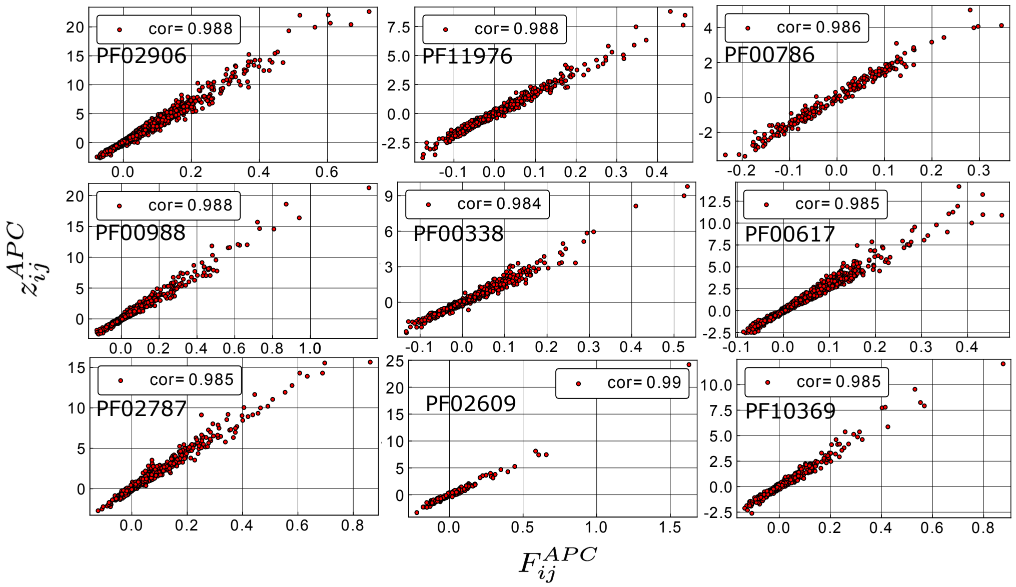

**Fig 13. z-scores of couplings derived from the natural MSA, as compared to the distribution of couplings derived from Null model III:** For each residue pair $(i, j)$, we calculate the z-score for the DCA score derived from natural data as compared to 50 realizations of Null model III. Results are shown for dataset DS1.
